# Epigenetically-regulated RNA-binding proteins signify malaria hypnozoite dormancy

**DOI:** 10.1101/2023.04.25.537952

**Authors:** Christa Geeke Toenhake, Annemarie Voorberg-van der Wel, Haoyu Wu, Abhishek Kanyal, Ivonne Geessina Nieuwenhuis, Nicole Maria van der Werff, Sam Otto Hofman, Anne-Marie Zeeman, Clemens Hendricus Martinus Kocken, Richárd Bártfai

## Abstract

**SUMMARY:** Dormancy enables relapsing malaria parasites, such as *Plasmodium vivax* and *cynomolgi*, to survive unfavorable conditions and maximize chances for transmission. It is caused by hypnozoites, parasites remaining quiescent inside hepatocytes before reactivating and establishing blood-stage infection. We integrated various omics approaches to explore gene-regulatory mechanisms underlying hypnozoite formation and reactivation. Genome-wide profiling of epigenetic marks identified a small set of genes that gets epigenetically silenced during hepatic infection of relapsing parasites. Furthermore, by combining single-cell transcriptomics, chromatin accessibility profiling and fluorescent *in situ* RNA hybridization, we show that these genes are exclusively expressed in hypnozoites and their silencing precedes parasite development. Intriguingly, these hypnozoite-specific genes mainly encode proteins with RNA-binding domains. We, hence, propose that repressive RNA-binding proteins keep hypnozoites in a developmentally competent but dormant state and heterochromatin-mediated silencing of the corresponding genes enables hypnozoite reactivation. Further testing of this hypothesis could provide clues for targeted reactivation and killing of these vicious pathogens.

## INTRODUCTION

Malaria parasites are unicellular eukaryotic pathogens from the *Plasmodium* genus. During their complex life cycle, they develop into various forms and adapt to vastly different environments within both the mammalian host and the mosquito vector (Cowman et al., 2016). While most of these developmental progressions are deterministic/unidirectional (e.g. asexual proliferation during blood stages or progression from the fertilized zygote to sporozoites within the mosquito), at two distinct points in the life cycle a ‘fork appears in the road’ and a cellular choice is made to maximize the chance of survival. The first and most well-known *Plasmodium* cell fate decision event is the commitment to gametocytogenesis, balancing the optimal rate between asexual proliferation in the mammalian host *vs* transmission through the mosquito vector to a new host. The other cellular detour is the less well characterized and more species-restricted ability to enter dormancy in the liver stage and become hypnozoites.

In the last decade, major advancements have been made in understanding the relevance of different gene regulatory mechanisms during life cycle progression. It has become clear that stage-specific ApiAP2 transcription factors are the main drivers of developmental progression (Modrzynska et al., 2017), while heterochromatin-mediated epigenetic regulation enables phenotypic variability and controls commitment to gametocytogenesis (Filarsky et al., 2018). Last but not least, translational repression of “pre-synthetized” mRNA transcripts in mature female gametocytes and sporozoites enables pausing and efficient continuation of the life cycle during host-to-vector and vector-to-host transitions (Lindner et al., 2019; Mair et al., 2006).

Despite these important discoveries, mechanisms controlling the second cell fate decision event, the formation and reactivation of relapsing malaria dormant liver stage forms remains a mystery (Schafer et al., 2021). Hypnozoites are not only a biological wonder, they are also a major challenge to *P. vivax* malaria control. These quiescent parasites can survive the current treatment regimens for blood stages and their reactivation results in relapse infections weeks to months later without renewed exposure to infectious mosquitos (Robinson et al., 2015). Although some seminal studies in the 1980s described the existence of persistent small liver stage forms (Krotoski, 1985), it took another four decades before it was finally demonstrated that these small liver stage forms can reactivate (Voorberg-van der Wel et al., 2020). Further underscoring the comparative paucity of knowledge, it was only very recently that the first insights into the transcriptome of hypnozoites were provided (Cubi et al., 2017; Gural et al., 2018; Mancio-Silva et al., 2022; Ruberto et al., 2022; Voorberg-van der Wel et al., 2017). These studies used different experimental approaches leading to substantial differences in the transcripts each study detected. Yet, there are a few important insights they all agree on: i) hypnozoites are transcriptionally active; ii) there are significantly fewer transcripts detected and hence less molecular functions supported in hypnozoites and iii) no gene with a strict hypnozoite-specific expression profile could be identified. Many of these studies identified a few ApiAP2 transcription factors with a putative role in regulating quiescence but their hypnozoite-specific expression, let alone function, has not been confirmed. Furthermore, the postulated role of epigenetic regulation in hypnozoite formation and reactivation has yet to be convincingly demonstrated (Dembele et al., 2014; Merrick, 2021).

## RESULTS & DISCUSSION

In line with the epigenetic processes that control gametocyte formation, we hypothesized that the control of liver stage development in relapsing malaria parasites might also have an epigenetic component. Therefore, to address if epigenetic changes do take place during liver stage development of relapsing parasites, we ChIP-seq profiled histone modifications associated with transcriptionally permissive euchromatic (H3K9/14ac) and silent heterochromatic state (H3K9me2) at stages preceding and following liver stage development of *P. cynomolgi*: sporozoites and blood stage, respectively. Upon visual investigation of the genome browser tracks (ChIP-over-input, Figure 1A) we observed generally broader heterochromatic domains in sporozoites as compared to blood stage parasites (blue rectangle on Figure 1A). However, we also detected a few small heterochromatic islands in blood stage parasites that were absent in sporozoites (red rectangle on Figures 1A, S1A,B). These epigenetic differences were confirmed by the opposite pattern in the H3K9/14ac ChIP-seq profiles (Figure 1A). By quantifying H3K9me2 ChIP-seq enrichment around the ATG of all *P. cynomolgi* genes (Figure 1B), we identified 404 genes in the broadened heterochromatic domains of sporozoites that were euchromatic in blood stage parasites (i.e. “heterochromatin losing”) and 25 genes that were exclusively heterochromatic at blood stages (i.e. “heterochromatin gaining”). Interestingly, by analyzing published H3K9me3/HP1 data (Fraschka et al., 2018; Vivax Sporozoite, 2019) from corresponding stages of *P. vivax* we identified a very similar set of largely homologous genes (322 heterochromatin losing and 9 gaining genes), demonstrating that these epigenetic changes are highly conserved amongst these relapsing parasites (Table S1).

**Figure 1.**
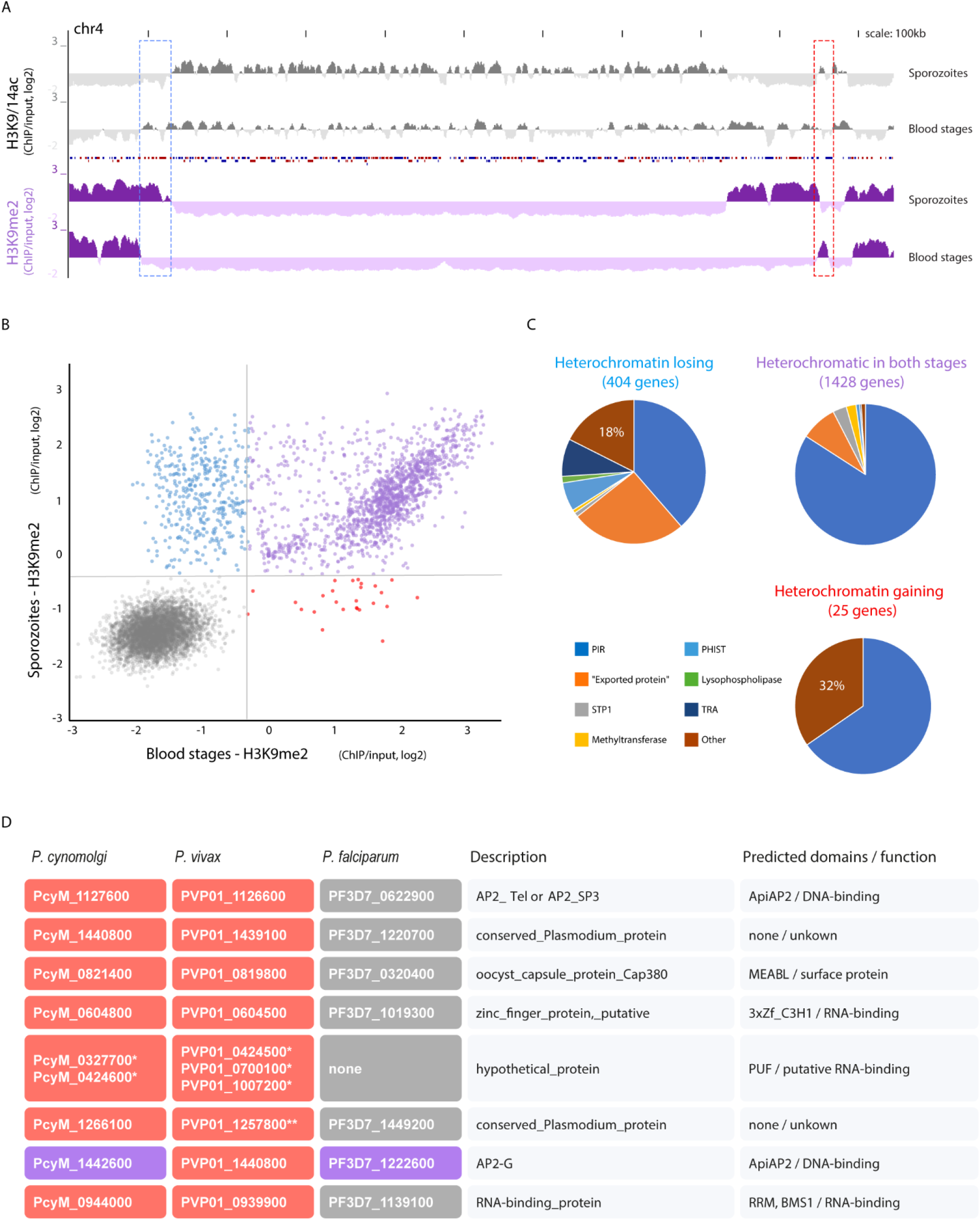
A sub-set of genes have alternate heterochromatin states during liver stage development in relapsing malaria parasites. **A)** Log2 transformed H3K9/14ac (grey) and H3K9me2 (purple) ChIP-over-input ratio tracks of chromosome 4 generated from *P. cynomolgi* salivary gland sporozoites and blood stage parasites. Two regions with opposing epigenetic changes are highlighted with red and blue rectangles, respectively. **B)** Scatter plot depicting H3K9me2 ChIP-over-input ratio for each *P. cynomolgi* gene in salivary gland sporozoites and blood stage parasites. Vertical and horizontal lines depict the cutoffs to call heterochromatic genes in either of the stages (probability > 0.99). Genes heterochromatic in both stages are colored purple, while euchromatic genes are grey. **C)** Pie-charts depicting the proportion of multigene family members among *P. cynomolgi* genes that are heterochromatic in sporozoites (blue), in blood stages (red) or in both (purple). Note the overrepresentation of non-multigene-family genes (“other”) among the heterochromatin gaining and losing categories. Similar data for *P. vivax* can be found in Table S1. **D)** Table of non-multigene-family genes that are heterochromatin gaining in *P. cynomolgi* and/or in *P. vivax* and their *P. falciparum* homologs. Genes are colored as in panel “**B**”. * homologues, non-synthenic group of genes with 2-3 copies in *P. cynomolgi* and *P*. vivax. ** based on visual inspection this gene also gains some heterochromatin, although not to the same extent as other heterochromatic genes (see Figure S1A).

As expected, most genes heterochromatic in both stages encoded exported proteins from various clonally variant multigene families (e.g. *pir, phist*, etc), while ‘single-copy’ genes were more abundant amongst heterochromatin losing (18%) and gaining (32%) genes (“other” on Figure 1C). Most heterochromatin losing single-copy genes with predicted function are involved in reticulocyte/red blood cell invasion or remodeling (e.g. *kahrp, rbp, clag*; Table S1), and hence it is conceivable that these genes need to lose heterochromatin and become activated prior to parasites commencing blood stage infection. On the other hand, single-copy heterochromatin gaining genes were enriched for proteins with predicted DNA- and RNA-binding properties, and they were heterochromatic at blood stages of both *P. cynomolgi* and *P. vivax* (Figures S1A,B). Importantly, the orthologs of these genes, except for AP2-G, appeared to be fully euchromatic in the non-relapsing malaria parasite, *P. falciparum* (Figure 1D). Amongst these genes were two ApiAP2 transcription factors: AP2-SP3 and AP2-G. AP2-SP3 has been found to be required for completion of sporozoite development in *P. berghei* (Modrzynska et al., 2017) and it remains possible that its function extends to the (early) liver stages. AP2-G, the master regulator of gametocytogenesis (Kafsack et al., 2014; Sinha et al., 2014), gains heterochromatin in *P. vivax* but to a much lesser extent in *P. cynomolgi* (Figures S1A,B). Interestingly, *de novo* mutation occurring in both of these factors have been found to display altered frequency in recurrent *P. vivax* infections (Dia et al., 2021) and they hence might be relevant for relapse. Furthermore, AP2-G has been recently shown to be expressed in a subset of *P. vivax* liver stage parasites (Mancio-Silva et al., 2022) and therefore its epigenetic regulation could contribute to early sexual commitment in this species. The list of heterochromatin gaining genes also contained two proteins with predicted RNA-binding properties (PcyM_0604800, PcyM_0944000) and four so far uncharacterized/hypothetical proteins. Upon closer investigation, we noticed that two of the latter (PcyM_0327700, PcyM_0424600) encode very similar proteins with putative pumilio homology domain (PUF, E-value: 2e-05 – 1e-10, Figure S1C). Importantly, such PUF proteins have been shown to function in translational repression in other stages of the parasite’s life-cycle (Miao et al., 2013; Muller et al., 2011). Collectively, these experiments revealed a small set of genes with gene-regulatory properties that gain heterochromatin *de novo* in relapsing malaria parasites as they transition through liver stages.

Next, we investigated whether these epigenetic differences also exist between day 6 hypnozoites and developing liver stage parasites using the chromatin accessibility assay (ATAC-seq). Using our unique dual-fluorescent *P. cynomolgi* reporter line (Voorberg-van der Wel et al., 2020) we sorted 1500-5000 developing liver stage, double-positive forms (mCherry^++^/GFP^++^) and an equal number of small liver stage forms (including hypnozoites) expressing lower amounts of GFP only (mCherry^-^ /GFP^+^) at 6 days post infection (Figure S2A).

Differential accessibility analysis of promoters of all protein coding genes by DiffBind (Ross-Innes et al., 2012) identified only 15 differential accessible promoters (FDR>0.05), suggesting that the overall chromatin/transcriptional landscapes of these two parasite populations are largely comparable (Figures 2A, Table S2). Interestingly, heterochromatin gaining genes with predicted RNA-binding properties all displayed significantly reduced accessibility in mCherry^++^/GFP^++^ parasites (Figures 2B,C; S2B; Table S2) while heterochromatin losing genes (blue) showed an opposing trend although their differential accessibility did not reach statistical significance (Figure 2B). It is possible that the mCherry^-^/GFP^+^ parasite population may not only contain ‘true’ hypnozoites but also some ‘retarded’ parasites and this potential mix could dampen the differences between the populations. With this caveat in mind, the differences that were still detected are highly suggestive that heterochromatin organization differs between hypnozoites and developing liver stage forms and that these epigenetic differences could contribute to hypnozoite-specific gene expression.

**Figure 2.**
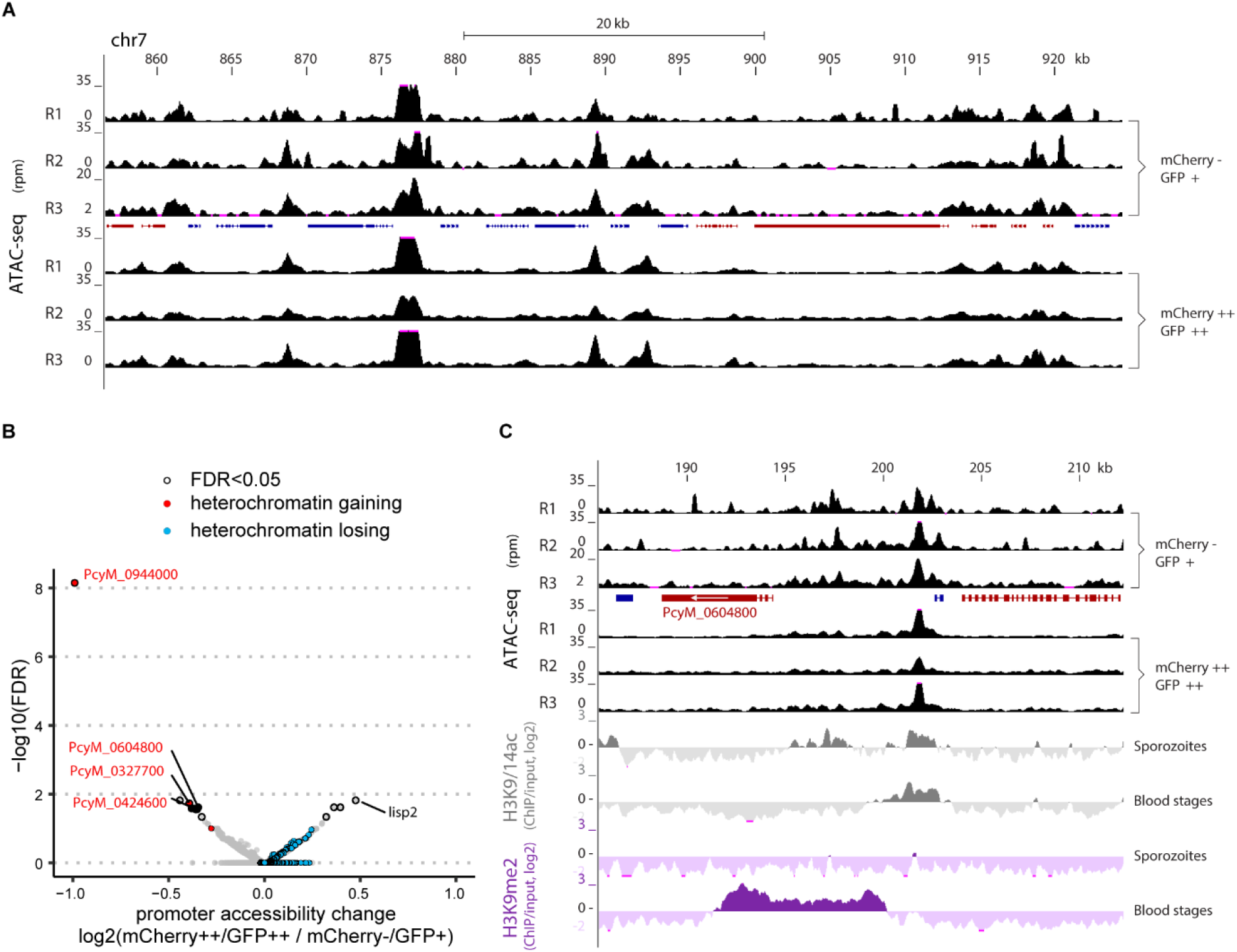
Chromatin accessibility is mostly similar in hypnozoites and liver schizonts with exception of some heterochromatin gaining and losing genes. **A)** ATAC-seq read per million coverage tracks obtained from three populations of mCherry^-^/GFP^+^ small liver stage forms (including hypnozoites) and mCherry^++^/GFP^++^ liver schizonts over a section of *P. cynomolgi* chromosome 7. **B)** Volcano plot of chromatin accessibility differences between mCherry^-^/GFP^+^ small liver stage forms and mCherry^++^/GFP^++^ liver schizonts in promoter region of *P. cynomolgi* genes. Promoters identified as significantly differentially accessible (FDR>0.05) are marked with black circles. Heterochromatic losing (blue) and gaining (red) genes are highlighted. **C)** ATAC-seq read per million coverage tracks of mCherry^-^/GFP^+^ small liver stage forms and mCherry^++^/GFP^++^ liver schizonts over the locus encoding for the heterochromatin gaining ZF-protein (PcyM_0604800). Log2 transformed H3K9/14ac (grey) and H3K9me2 (purple) ChIP-over-input ratio tracks from Figure 1 are also shown. “Blue genes” are transcribed from left to right, while “red genes” from right to left. For other examples see Figure S2B.

To test this hypothesis and investigate gene expression changes associated with parasite development in general, we performed single-cell RNA-seq analysis of flow cytometry-sorted, infected hepatocytes at 2, 3, 4 and 6 days post sporozoite inoculation from three independent cultures (Figure 3). With the cut-off for positive detection set at a minimum of 400 genes per cell with each gene being detected in at least five cells per batch, we obtained a dataset of 1243 cells and 4773 genes. Parasite cell clustering was largely driven by the time after infection (Figure 3A) and associated differences in steady-state mRNA abundance (Figure S3). Most day 2 parasites (Figure 3A, top panel, light blue cells) clustered on the left side of the UMAP, while the most developed mCherry^++^/GFP^++^ parasites localized to the top right (Figure 3A, middle panel) with a gradient of day 3-4 parasites in between (Figure 3A). However, there is a distinct subset of day 6 parasites (grey dots at the lower half of the UMAP) that did not cluster with these well-developed forms and appeared to display transcript abundances similar to day 3-4 parasites (i.e. stalled in development). Accordingly, unsupervised clustering of these cells yielded 5 parasite clusters where cluster 1 mainly contained day 2 parasites, clusters 4 and 5 represented the most developed liver stage parasites, and clusters 2 and 3 contained in-between liver stage forms, which would likely include stalled parasites and/or candidate hypnozoites (Figure 3A; lower panel). In parasite cluster 4, and in particular cluster 5, nearly all genes displayed higher mRNA abundance and a much higher total amounts of steady-state mRNA (Figure 3B), as observed earlier in bulk RNA-seq (Voorberg-van der Wel et al., 2017). This observation that differences between parasite clusters are rather quantitative than qualitative, together with the overall similar chromatin accessibility pattern of small and developing forms (Figure 2), suggests that the overall transcriptional profile of hypnozoites and liver schizonts is somewhat surprisingly comparable and hence transcriptional changes might not be the main drivers of hypnozoite dormancy/reactivation.

**Figure 3.**
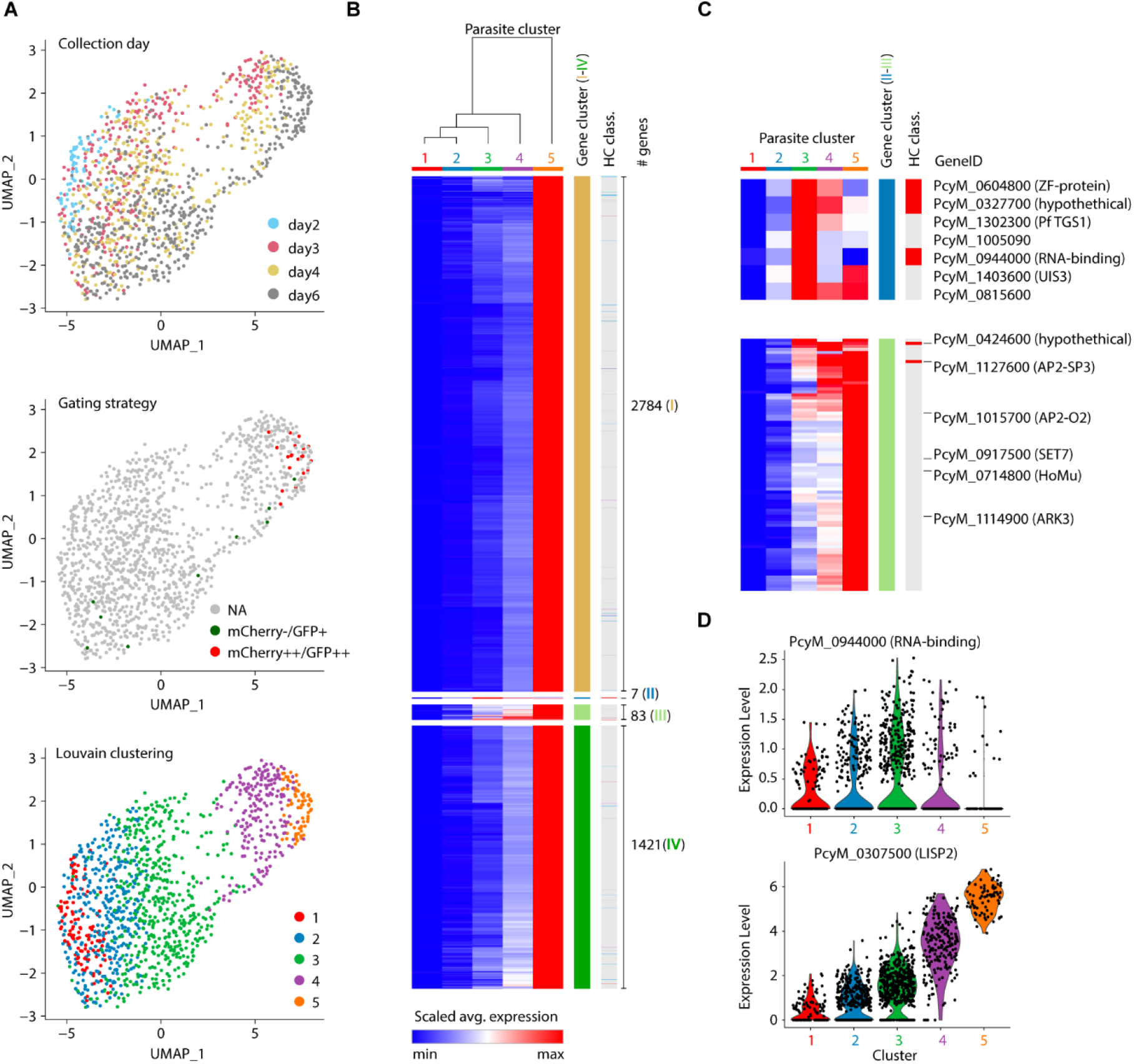
Single-cell RNA-seq analysis of *P. cynomolgi* liver-stage parasites reveals differential expression of heterochromatin gaining genes in early liver stage forms. **A)** Uniform Manifold Approximation and Projection (UMAP) of 1243 liver stage *P. cynomolgi* parasites based on their gene expression profile. Top panel: colored based on age of the parasites (days post infection); middle panel: twenty index-sorted day 6 mCherry^-^/GFP^+^ (green) and mCherry^++^/GFP^++^ (red) parasites are highlighted; bottom panel: colored based on unsupervised clustering of expression profiles (Louvain clustering at resolution 0.4 of PCs 1-15). For further details see Figure S3. **B)** Heatmap with relative gene expression profiles of 4295 *P. cynomolgi* genes across the different parasite clusters (1-5) as in panel **A**. Within gene clusters (I-IV), rows were ordered by hierarchical clustering (Euclidean, average linkage). **C)** Zoom in of heatmap of gene clusters II and III from panel **B**. Name and gene ID of some genes of interest as well as heterochromatin classification (HC class.) from **Figure 1. B** is indicated. **D)** Violin plots of expression levels of a heterochromatin gaining gene from gene cluster II (PcyM-0944000, RNA-binding protein) and a well know marker of early liver-stage development (PcyM-0307500, *lisp2 (Gupta et al*., *2019)*) (see also Table S3).

To identify genes with relatively higher expression in earlier stages, we clustered the average gene expression per parasite cluster using k-means clustering. This yielded four gene clusters (I-IV, Figure 3B, Table S3). The two smaller gene clusters contained genes whose average expression per parasite cluster deviated from the overall trend (Figures 3B-C, gene clusters represented by blue (II) and light green (III) bars on the right side of the heat maps). Particularly, the seven genes belonging to gene cluster II showed high expression in cluster 3 parasites and an analysis of variance using the Kruskal-Wallis rank sum test yielded significant differences in mean expression among parasite clusters for all these genes except for PcyM_0815600 (adjusted p-value < 0.05, Table S4). Furthermore, post-hoc Dunn’s tests showed that expression levels for PcyM_0604800, PcyM_0944000, PcyM_1302300, PcyM_1005090 and PcyM_1403600 were significantly higher between parasite cluster 3 and parasite cluster 4 or 5 (adjusted p-value < 0.05, Table S4). Notably, nearly all heterochromatin gaining single-copy genes and in particular the ones with predicted RNA-binding properties (PcyM_0604800, PcyM_0944000, PcyM_0327700, PcyM_0424600) belonged to these gene clusters. Furthermore, gene cluster II contained one additional gene (PcyM_1302300; *tgs1*, trimethylguanosine synthase) that has a predicted function in mRNA processing (Bawankar et al., 2010). Gene cluster III, on the other hand, contained 83 genes with overall higher expression levels in the parasites of parasite clusters 3 and 4, as compared to most other genes. However, all genes with significant differences in gene expression across parasite clusters (Kruskal-Wallis rank sum test, adjusted p-value < 0.05) showed higher expression in parasite cluster 5 compared to parasite cluster 3 or 4 (Dunn’s Test, see output in Table S3). Interestingly, this gene cluster contained two ApiAP2 transcription factors: the heterochromatin gaining AP2-SP3 (PcyM_1127600) and AP2-O2 (PcyM_1015700), which might be required for establishment of the early liver stage specific gene expression program. Of note, we did not detect significant levels of transcript for AP2-G (PcyM_1442600) in *P. cynomolgi* liver stage parasites.

Next, we set out to validate the expression patterns of some of these RNA- and DNA-binding proteins with fluorescence *in situ* hybridization (RNA-FISH). For two of the heterochromatin gaining RNA-binding proteins (PcyM_0604800 and PcyM_0944000) we performed FISH at days 2, 3, 4 and 6 of liver stage development (Figure 4, Table S4). While the control probe against the housekeeping gene *gapdh* (PcyM_1250000, green) detected both small uninucleate forms (1n) as well as multinucleate (>1n) liver schizonts, signals obtained with probes against both RNA-binding protein mRNAs (red) were largely restricted to the small forms at all time-points (Figure 4A). In fact, while we detected FISH signal in most (70-100%) uninucleate forms, only a few multinucleate liver schizonts at day 4 and 6 displayed minimal levels of signal (Figure 4B).

**Figure 4.**
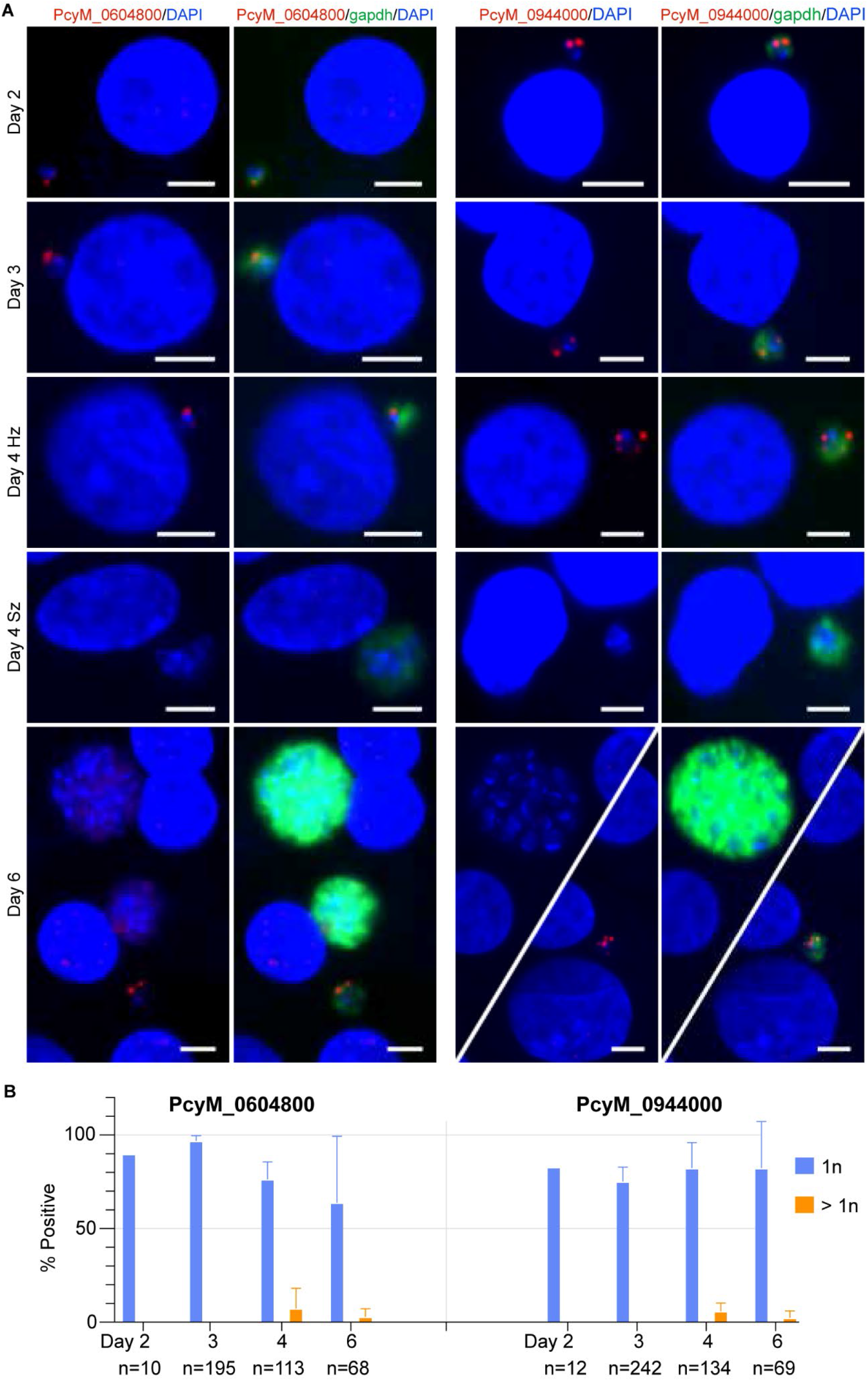
FISH analysis of two heterochromatin gaining genes validates hypnozoite-specific expression and silencing in developing parasites. **A)** Fluorescence *in situ* RNA hybridization images of heterochromatin gaining, gene cluster II RNA-binding protein gene transcripts (PcyM_0604800 and PcyM0944000, red) and the house-keeping gene transcript (PcyM_1250000 / *gapdh*, green) in *P. cynomolgi* parasite-infected hepatocyte cultures at different time-points after infection (day 2-6). DNA of the parasites as well as host hepatocytes is visualized by DAPI staining (blue). Scale bar: 5 µm **B)** Bar graph depicting the percentage of uninucleate (1n) and multinucleate (>1n) liver stage *P. cynomolgi* liver stage parasites with FISH signal using probes against PcyM_0604800 and PcyM0944000 respectively at different days post infection. Error bars depict standard deviation calculated based on multiple independent experiments (Table S4).

FISH results of two more genes from gene cluster II (PcyM_1005090 and PcyM_1302300) similarly showed more pronounced transcript abundance in hypnozoites compared to schizonts albeit quantification was less extensive (Figure S4, Table S4). In contrast, mRNA of the two ApiAP2 transcription factors in gene cluster III were detectable both in uninucleate and multinucleate liver stage forms (Figure S4). Collectively, these FISH experiments fully validate our single cell RNA-seq data and further strengthen the strictly small-form-specific transcription of three putative RNA-binding proteins (PcyM_0604800, PcyM_0944000 and PcyM_1302300).

In summary, via multi-omics profiling we have identified a small set of marker genes for hypnozoites (see Table S5 for overview). The expression of these epigenetically-regulated RNA-binding proteins inversely correlates with progression of liver stage development in a relapsing malaria parasite. While the exact function of these proteins remains to be determined, given the prominent role of translational repression at other stages of the parasite’s lifecycle (Mair et al., 2006; Turque et al., 2016), we hypothesize that these RNA-binding proteins are also involved in a similar process (i.e. translational repression or delayed processing of mRNAs required for liver stage proliferation). In fact, sporozoite maturation and transmission to the human host involve waves of translational repression that are regulated by RNA-binding proteins (Gomes-Santos et al., 2011; Lindner et al., 2019; Muller et al., 2011; Muller et al., 2021). Accordingly, RNA-binding proteins identified in this study could have evolved to facilitate a “natural extension” of these translational repression events during early liver stages. In line with this hypothesis a recent study, has also identified RNA-binding proteins to be differentially expressed between hypnozoites and liver schizonts in *P. vivax* (Ruberto et al., 2022). Notably, however, the set of RNA-binding proteins identified in this study is largely distinct from the one identified by Ruberto et al. (except for PVP1_0939900) and hence it remains to be determined if and how these different translational regulatory processes cooperate in regulating cell fate of liver stages of relapsing malaria parasites. Importantly, in none of the previously identified translational repressor processes, epigenetic silencing of translational repressors was observed. Hence, it seems plausible that similar to epigenetic regulation of the master regulator of gametocytogenesis (AP2-G(Coleman et al., 2014; Filarsky et al., 2018; Kafsack et al., 2014; Sinha et al., 2014)), heterochromatin-mediated silencing of the hypnozoite-specific RNA-binding protein genes identified here could constitute an epigenetic switch controlling the fate of infection by alleviating developmental repression (model under Supplemental Information). Further exploration of this hypothesis could resolve the mystery of relapsing malaria parasites and instruct strategies for targeted reactivation and killing of these important human pathogens.

## ACKNOWLEDGEMENTS

We are thankful to F. van Hassel for preparing graphical representations and to Nicholas Proellochs for proofreading and editing of the manuscript. We are also grateful to colleagues at the mosquito breeding facility of Radboudumc (Nijmegen) for provision of *Anopheles stephensi* mosquitoes and colleagues at the Animal Science Department (BPRC, Rijswijk) for animal care and veterinary assistance. We thank Herman Oostermeijer for technical support in flow cytometry sorting experiments. This project greatly benefited from the resources and services provided by PlasmoDB (plasmodb.org). We would specifically like to acknowledge the help of Mark John Hickman with identification of orthologous genes between *P cynomolgi* and *vivax*. This project obtained financial support from the Dutch Research Council (ZonMW-TOP-grant #91218010).

## AUTHOR CONTRIBUTIONS

**C.G.T**. and **A.V-vdW**. designed and performed experiments, analyzed and interpreted data, prepared illustrations and wrote parts of the manuscript. **H.W**. performed analysis of ATAC-seq data and prepared illustrations. **A.K**. performed bioinformatic analysis of the ChIP-seq data. **I.G.N**., **N.M.vdW**. and **A-M.Z**. contributed with production of blood stage parasites and sporozoites and liver stage cultures. **S.O.H**. performed flow cytometry sorting and analysis, prepared relevant illustration and wrote corresponding parts of the manuscript. **C.H.M.K**. and **R.B**. conceived the study, designed and supervised experiments and provided resources. **R.B**. furthermore performed ChIP-seq and ATAC-seq experiments and wrote the manuscript. All authors contributed to the editing of the manuscript.

## METHODS

### Lead contact

Further information and request for datasets or other resources should be directed to the lead contact, Dr Richard Bartfai.

### Data availability

Raw and processed next generation sequencing datasets have been submitted to the NCBI Gene Expression Omnibus and are accessible via reference number GSE180258. The corresponding subseries numbers are: GSE180256 (ChIP-seq), GSE180255 (ATAC-seq) and GSE180257 (scRNA-seq).

### Ethics statement

*P. cynomolgi* parasites and hepatocytes were obtained from nonhuman primates because no other models (*in vitro* or *in vivo*) were suitable for this project. All experiments were performed according to Dutch and European laws. BPRC has been fully accredited by the Council of the Association for Assessment and Accreditation of Laboratory Animal Care (AAALAC International). The work was performed under ‘Centrale Commissie Dierproeven’ CCD license number AVD5020020172664, and the research protocol was approved by the local independent ethical committee conform Dutch law (BPRC Instantie voor Dierenwelzijn, IvD, agreement number # 007). The rhesus monkeys (*Macaca mulatta*, male, age 7-15 years, Indian origin) used for this research were captive-bred and socially housed. Housing, feeding and environmental enrichment was as described previously (Voorberg-van der Wel et al., 2017).

### Transgenic *Plasmodium cynomolgi* sporozoite production

Parasite donor monkeys were infected with a dual fluorescent *P. cynomolgi* M line that expresses mCherry under control of liver schizont specific *lisp2* (PcyM_0307500) promoter and 3’ UTR regions and GFP (and selectable marker) under control of the constitutive *hsp70* promoter (PcyM_0515400) (Voorberg-van der Wel et al., 2020). A centromere present in the construct ensures that the line is maintained throughout the life cycle (Voorberg-van der Wel et al., 2013). Parasite infections and feedings of mosquitoes (two to five days old female *Anopheles stephensi* mosquitoes Sind-Kasur strain; Nijmegen RadboudUMC, Department of Medical Microbiology) were performed as previously described (Voorberg-van der Wel et al., 2017). Two weeks post mosquito feeding, salivary gland sporozoites were isolated for hepatocyte infection.

### *P. cynomolgi* liver stage culture

Primary hepatocytes from *Macaca mulatta* or *Macaca fascicularis* were isolated freshly or thawed from frozen stocks as described previously (Zeeman et al., 2016). Hepatocytes were plated into collagen coated 96-well plates (65 × 10^3^ hepatocytes per well; Perkin Elmer) and maintained at 5% CO_2_ at 37 °C in William’s B medium supplemented with 2% DMSO: William’s E medium with glutamax containing 10% human serum (A+), 1% MEM non-essential amino acids, 2% penicillin/streptomycin, 1% insulin/transferrin/selenium, 1% sodium pyruvate, 50 µM β-mercapto-ethanol, and 0.05 µM hydrocortisone. After 2-3 days, hepatocytes were inoculated with 50 × 10^3^ *P. cynomolgi* sporozoites per well. Cultures were maintained at 5% CO_2_ at 37 °C with 2-3 medium refreshments (William’s B without DMSO) per week until flow cytometry sorting or fixation. Quality of the cultures was assessed by Operetta high content imaging as described previously (Voorberg-van der Wel et al., 2021; Voorberg-van der Wel et al., 2022).

### Flow cytometry cell sorting

At days 2, 3, 4 and 6 post-sporozoite inoculation, liver stage cultures were harvested by trypsinization (3-5 min. at 37 °C (Voorberg-van der Wel et al., 2021)). Samples were washed, passed through a 100 µm cell strainer, resuspended into 20% William’s B medium and kept on ice until sorting. Sorting was performed in one experiment with a 3-laser, 8-color FACSMelody (BD Biosciences) and for all other experiments a 4-laser, 16-color FACSAriaIIIu (BD Biosciences) was used. GFP was excited by 488 nm laser and emission was collected through a 527/32 filter (FACSMelody) or a 530/30 filter (FACSAria). mCherry was excited by a 561 nm laser, and emission signals were obtained through a 613/18 filter (FACSMelody) or a 610/20 filter (FACSAria). The general performance of the FACSAria was checked by C(ytometer)S(etup) &T(tracking) beads. A N(oise) D(ensity) filter 1.5 was placed in front of the FSC diode to reduce the signal. Additionally, Cyto-cal multifluor beads (Thermo Scientific; #FC3MV) were used to manually adjust voltage settings to a fixed Median Fluorescence Intensity (MFI; FITC=100, ECD=102) to ensure the same target fluorescence intensity. Samples were acquired at a flow rate (1-4) corresponding to an event rate of approximate 5000 events/sec. Sorting was performed using a 100 µm nozzle at 20 PSI or with a 130 µm nozzle at 11.5 PSI. The drop delay was set automatically before the experiment by the Autodelay wizard using BD FACSTM Accudrop Beads (BD Biosciences). GFP positive parasites (see below) were collected in 384 wells plates containing 100 nl lysis solution with the sorting precision at “single cell” (yield mask: 0; purity mask: 32; phase mask: 16) to maximize the purity and counting accuracy. By including the index sorting function, information on parameters such as forward scatter (FSC), side scatter (SSC), GFP and mCherry signals was recorded for each well containing a sorted event. In this way, information could be traced back to specific wells of interest. As controls, single cell sorts were performed through gating on uninfected hepatocytes, mCherry^++^/GFP^++^parasites (schizonts) and mCherry^-^/GFP^+^ parasites (hypnozoites) (Figure S2A for an example and (Voorberg-van der Wel et al., 2020)). Immediately after sorting, plates were centrifuged (2 min., 350 g, 4 °C) and stored at -80 °C until further processing.

### ChIP-seq

For ChIP-seq of *P. cynomolgi* M blood stage parasites, on two sequential days, 9 ml heparin blood was collected from a donor monkey (containing 1.2 % ring stage parasites and 1.4% trophozoites respectively). For ChIP-seq of *P. cynomolgi* M sporozoites, 6.5 × 10^6^ salivary gland sporozoites were isolated two weeks post-mosquito-feeding, collected in PBS/glycerol and stored frozen until further processing.

Native ChIP was carried out as described earlier (Hoeijmakers et al., 2012). In short: After saponin lysis of the red blood cells and/or lysis of the parasites, nuclei were separated using a 0.25 M sucrose buffer cushion. Native chromatin was prepared by MNase digestion and subsequent extraction with salt-free buffers (10 mM Tris pH 7.4, 1 mM EDTA; 1 mM Tris pH 7.4, 0.2 mM EDTA). Chromatin was diluted in 2×ChIP incubation buffer (100 mM NaCl, 20 mM Tris pH 7.4, 6 mM EDTA, 1% Triton X-100, 0.1% SDS). ∼400 ng DNA-containing chromatin was incubated with 1 µg antibody (H3K9/14ac (Diagenode, C15410005); H3K9me2 (in house, corresponds to Merck Millipore 07-441) overnight at 4°C followed by the addition of 10 µl A/G beads (SantaCruz Biotechnology) and further incubation for 2 h. After washing with buffers containing 100, 150 and 250 mM NaCl, immuno-precipitated DNA was eluted and purified using PCR purification columns (Qiagen). 1-10 ng of ChIP-ed or input DNA were used to generate sequencing libraries. All samples were first end-repaired, a 3’ A-overhang added and NextFlex 6bp barcoded adapters (Bio Scientific) were ligated. Adapter ligation was followed by 2 successive Agencourt AMPure XP bead purifications (Beckman Coulter) and library amplification by a Plasmodium-optimized KAPA PCR amplication protocol. First, 2x KAPA HiFi HotStart ready-mix (KAPA Biosystems) and NextFlex PCR primers were used for 4 cycles of PCR amplification (98 °C for 2 min; 4 cycles of 98 °C for 20 sec, 62 °C for 3 min; 62 °C for 5 min). Subsequently, amplified libraries were size selected for ∼270 bp (mono-nucleosomes + ∼125 bp NextFlex adapter) using 2% E-Gel Size Select Agarose Gels (Thermo Fisher Scientific) and again amplified as described above for 9 cycles. To deplete adapter dimers and clean-up the DNA, libraries were subsequently purified with Agencourt AMPure XP bead (Beckman Coulter) (1:1 library to bead ratio) before sequencing.

ChIP-seq and input libraries were sequenced on the Illumina NextSeq 500 system with a ∼20% phiX spike-in (Illumina, FC-110-3001) to generate 75 bp single-end reads (NextSeq 500/550 High Output v2 kit). The quality of the resulting reads was checked with FastQC (V0.11.8) and the reads were mapped against the P. cynomolgi reference genome from PlasmoDB (www.plasmodb.org) using BWA samse (v0.7.17-r1188). Mapped reads originating from the mitochondrial and apicoplast genome, multi-mapping reads, and reads having a mapping quality below 15 were removed (SAMtools v1.9). Before visualizing the ChIP-seq data in the UCSC Genome browser, the bam file for each library was normalized to its respective sequencing-depth, quantified as counts per million, with the deepTools (ver. 3.5.1) bamCoverage command. The resulting sequencing-depth normalized coverage files (obtained in bigwig format) were used to calculate the ChIP enrichment over the input sample by dividing ChIP sample files with input sample files with a subsequent log2 transformed (one pseudo count was added to avoid division by zero). This operation was performed with the bigwigCompare command in the deepTools package (ver. 3.5.1). For visualization within the UCSC genome browser, tracks were smoothened (4) and the windowing function was set as ‘mean’.

In order to obtain a per gene H3K9me2 enrichment values, log2 ChIP-over-input ratios were calculated in windows ±500bp in relation to the ATG of all protein coding genes in the sporozoite and blood stage of *P. cynomolgi* or *P. vivax* genomes. The bigwigCompare command from deepTools package was implemented for this purpose using a custom generated BED file with +/-500bp regions flanking ATG as reference. To identify and compare the sets of heterochromatic genes in sporozoites and blood stages respectively ChIP-seq enrichment values were assigned to either a ‘heterochromatic’ or ‘euchromatic’ compartment using a bivariate Gaussian mixture model (genes with p > 0.99 in the second compartment were considered heterochromatic). The normalmixEM algorithm from the mixtools package (ver. 1.2.0) in R was used for this purpose. Genes were categorized as heterochromatin gaining or losing if they fell into the heterochromatin compartment in one but not the other stage.

### ATAC-seq

For ATAC experiments, three bulk populations of 1500-5000 day 6 hypnozoite-(mCherry^-^/ GFP^+^) and schizont-(mCherry^++^/GFP^++^ infected hepatocytes were collected in two independent experiments. Sorting was performed using the AriaIIIu as described previously (Voorberg-van der Wel et al., 2021). Parasites were collected into PBS containing 1% BSA, centrifuged (5 min, 500g, 4 °C) and kept on ice until transposase treatment. Transposase treatment was performed in 10ul volume containing 1xTD buffer (Nextera kit, Illumina), 0.5ul “home-made” Tn5 transposase adapter complex and 0.4mg/ml digitonin (Promega, #G9441) at 30 min at 37 °C. After column purification (MinElute PCR Purification Kit, Qiagen) transposed DNA fragments were PCR amplified (98 °C for 2 min; 13 cycles of 98 °C for 20 sec, 62 °C for 3 min; 62 °C for 5 min) using Nextera i5/i7 indexing primers (Illumina) and 2x KAPA HiFi HotStart ready-mix (KAPA Biosystems). The resulting libraries were subsequently purified with Agencourt AMPure XP bead (Beckman Coulter) first at 1:0.65 ratio to remove large DNA fragments, followed by addition of extra beads to the supernatant to 1:1.8 ratio and removal of adapter dimers. ATAC-seq libraries were sequenced on the Illumina NextSeq 500 system (Illumina, FC-110-3001) to generate 2x38 bp paired-end reads with the corresponding indexes.

For ATAC-seq data analysis, pair-end reads were mapped to a merged genome reference of Mmul_10 (https://www.ensembl.org/Macaca_mulatta/Info/Index) and PlasmoDB44 (https://plasmodb.org/plasmo/app/downloads/release-44/PcynomolgiM/) using BWA 0.7.17-r1188 mem (Li and Durbin, 2009) with default settings. Next, reads mapped to the PlasmoDB44 genome only were extracted for further analysis. Duplicates and reads with mapping quality lower than 30 were removed using samtools 1.7 (Li et al., 2009). Open chromatin profile tracks were generated using deeptools 3.5.0 bamCoverage function (Ramirez et al., 2016) with the following setting, --normalizeUsing CPM. To define the promoter regions a part of the intergenic region upstream of the ATG of each gene was selected. In case of intergenic regions between two genes with the same orientation (“tandem”), two third of intergenic region upstream of the ATG was assign to the corresponding gene as its promoter. For convergent genes, half of the intergenic region was assigned to one gene while the other half to another gene as promoters. Counts of all replicates in promoter regions of protein coding genes (excluding those on mitochondrial and apicoplast contigs) were processed in DiffBind (v3.4.0) (Ross-Innes et al., 2012) with default settings except that counts were normalized by reads in promoter regions.

### Single-cell RNA sequencing

The single-cell RNA-seq libraries were prepared following the CEL-seq2/SORT-seq protocol (Hashimshony et al., 2016) with CEL-seq2 primer design as in (Gerlach et al., 2019). In short, single cells were sorted based on their GFP signal into 384-well plates containing 100 nl of unique CEL-seq2 compatible primer (7 ng/µl) and 5 µl mineral oil (Sigma-Aldrich). Following cell lysis and denaturation, reverse transcription was performed using 3.5 units SuperScript II reverse transcriptase per well (Invitrogen, #18064014) in the presence of 20 nl ERCC spike-in per well (Ambion™ 4456740, 1:5000 diluted in H_2_O).

Single-cell RNA-seq libraries were sequenced for 2x 42 bp on the Illumina NextSeq 500 system (read 1 is part of the 3’ end mRNA, read 2 contains the UMI and cell barcode). Sample demultiplexing based on the cell barcode was performed with UMI-tools v1.0.0 using the extract command which append mCherry^++^/GFP^++^s the UMI from read 2 to the ID of read 1. Demultiplexed read 1 fastq files were processed with Trim Galore v0.6.4 with default settings to remove adapters, low-quality bases, polyA and polyT repeats. PolyG sequences were removed using fastp v0.20.0(Chen et al., 2018) (these polyG sequences are an apparent artefact of NEXT500 sequencing). Demultiplexed and trimmed read 1 fastq files were mapped with STAR 2.4.0j with default settings to a reference genome composed of the *P. cynomolgi* M PlasmoDB release 44 genome, the sequence of the *mcherry-gfp* transgene with 33 bp 3’UTR on both sides, the NCBI M mulatta v10 reference genome and the control ERCCs. Bam files were filtered for a mapq of 255 and the readID was corrected with a custom script to remove the tags added in the polyA- and polyT trimming procedure by Trim Galore, this step was required such that UMI-tools could recognize the UMI again. Bam files were split per reference genome and these were then used by featureCounts v1.6.4 (Subread package, http://subread.sourceforge.net/) to assign reads to genes with settings –s 1 for strand-specific counting in forward mode and –R bam to get output in bam file. Since 3’ UTRs are not annotated in the *P. cynomolgi* reference genome, we extended the 3’ end of each gene in PlasmoDB-44_PcynomolgiM.gff with 1kb, or less when this 1kb overlapped with genes on the same strand. After read-to-gene assignment, UMIs were counted with the UMI-tools count command with settings --per-gene --gene-tag=XT --per-cell and --extract-umi-method=read_id.

Downstream data analyses were performed in R. Cells from the three batches were treated separately until integration. For each batch, cells with gene counts from *P. cynomolgi, M. mulatta* and ERCC were imported into Seurat (Seurat package v3.1.5 (Hao et al., 2021)) and filtered for at least 400 *P. cynomolgi* genes. Candidate duplicate events were removed with a *P. cynomolgi* UMI count filter of 40.000. Genes were filtered on being detected in at least five cells per batch, resulting in a dataset of 1243 cells and 4773 genes.

*P. cynomolgi* gene counts were normalized per batch using the “Normalization by spike-ins” strategy, as described in the online companion of Amezquita *et al (Amezquita et al*., *2020)* . To do so, filtered counts in Seurat objects were converted to SingleCellExperiment objects and with splitAltExps (SingleCellExperiment package v1.8.0 (Amezquita et al., 2020)) ERCC and *M. mulatta* counts were placed in different alternative experiments. The ERCC alternative experiment was used by computeSpikeFactors (scran package v1.14.6 (Lun et al., 2016)) to compute the scaling factors to be used to normalize the counts with in the logNormcounts (scater package v1.14.6 (McCarthy et al., 2017)). This normalization was performed with and without log-transformation per batch (not across batches since the added amount of ERCC to the reaction mixture is more equal within than between batches). The ERCC-normalized counts were imported into the “RNA” assay of a new Seurat object: ERCC-corrected counts in the “counts” slot, the log-ERCC-corrected counts in the “data” slot.

The default integration workflow of Seurat was followed to integrate the counts from the three experiments. Integration was performed based on features detected by FindVariableFeatures with “vst” method and for 2000 features. Anchor finding was performed with the FindIntegrationAnchors function with default settings except for the k.filter which was set to 100 anchors. Anchor finding and integration used dims 1:30. The integrated data was scaled, PCA dimensionality reduction was run and UMAP and Louvain clustering were performed following the standard Seurat integration workflow. UMAP and clustering used PCs 1 till 15. Louvain clustering resolution was set to 0.4 resulting in 5 parasite clusters.

The dotplot function from Seurat was used to obtain average and scaled average expression per parasite cluster from the RNA assay of the integrated Seurat object (ERCC-scaled counts). For the heatmap of scaled average expression, genes within the lowest decile (10%) of the summed average expression were removed leaving 4295 genes. The heatmap was generated with the Morpheus tool of the broad institute (https://software.broadinstitute.org/morpheus/). Using the scaled data from the dotplot function. Coloring was set as relative to row minimum and row maximum. Rows (genes) were clustered by k-means clustering using 1-pearson correlation with 1000 iterations. Within gene clusters, rows were ordered by hierarchical clustering (Euclidean, average linkage).

To determine expression level differences between parasite clusters for genes in gene cluster II and gene cluster III, the normalized gene counts were extracted from the Seurat object using FetchData and a Kruskal-Wallis rank sum test was performed to test for differences of the medians between parasite clusters (1 – 5) using the Kruskal().test function in R. p-values were adjusted for multiple comparisons using the p.adjust() function using the Benjamini & Hochberg correction methods. For the genes that showed and adjusted p value < 0.05, a Dunn’s test was performed using the dunnTest function from the FSA package (Ogle et al., 2021), and in one-sided mode. Also, for the Dunn’s test the Benjamini & Hochberg method was used to prevent inflation of Type I errors.

### RNAscope *in situ* hybridization

*P. cynomolgi* M infected primary macaque hepatocytes were cultured for 2, 3, 4 or 6 days in collagen coated CellCarrier-96 well plates (Perkin-Elmer) followed by fixation for 30 min at room temperature in 4% paraformaldehyde in PBS (Affymetrix). The material was dehydrated and stored in 100% ethanol at -20 °C until further processing. RNA *in situ* detection was performed using the RNAscope Multiplex Fluorescent V2 Assay kit (Advanced Cell Diagnostics) in combination with OPAL dyes 520 and 570 (Akoya Biosciences) according to the manufacturer’s instructions. RNAscope probes used are shown in Table S6. Images were obtained using a Leica DMI6000B inverted fluorescence microscope equipped with a HC PL APO 63x/1.40-0.60 oil objective and with a DFC365FX camera.

