## Supplementary data for "Epigenetically-regulated RNA-binding proteins signify malaria hypnozoite dormancy"

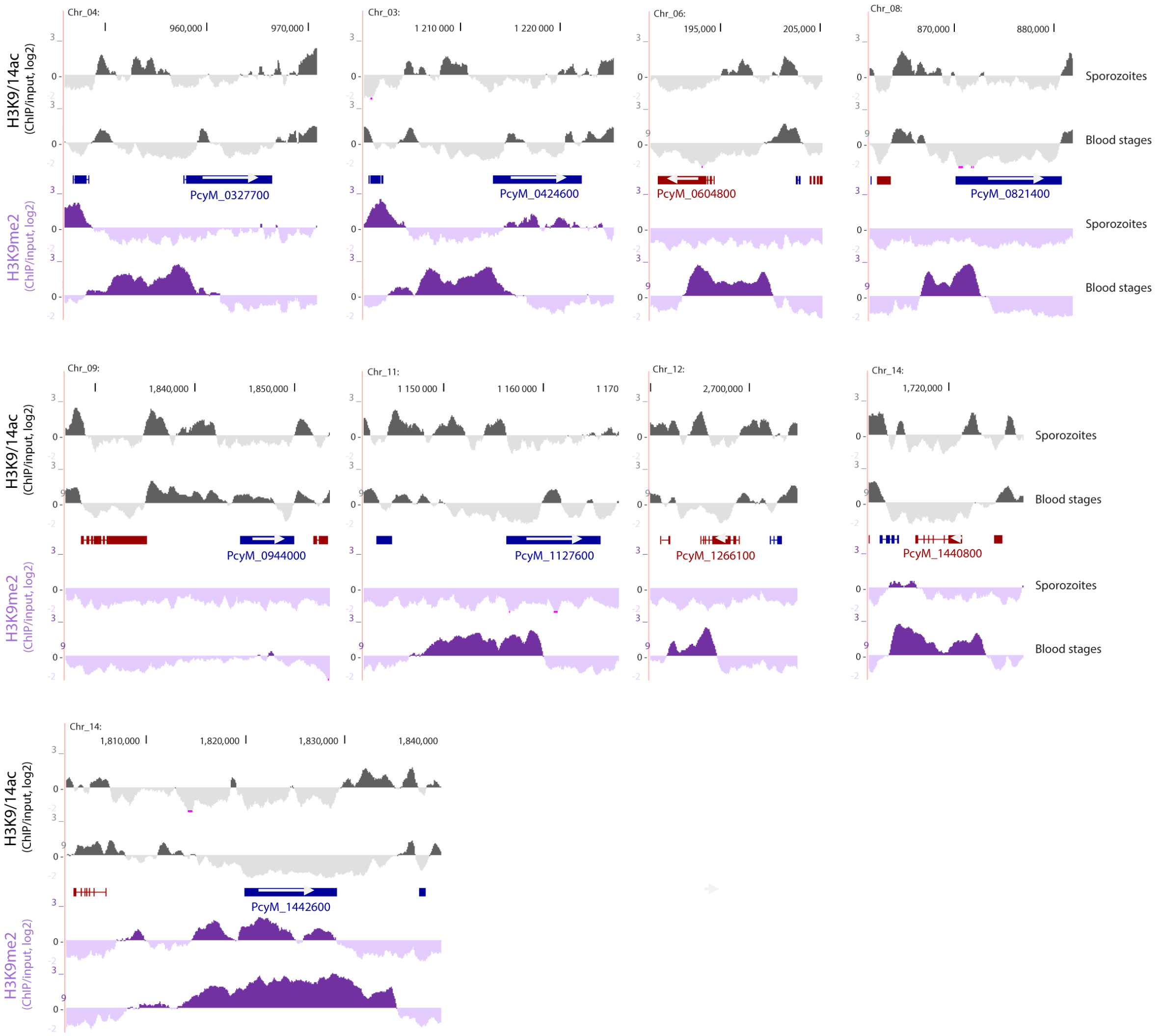


**Figure S1A: Epigenetic profile of *P cynomolgi* genes listed on Fig. 1d**

Log2 transformed H3K9/14ac (grey) and H3K9me2 (purple) ChIP-over-input ratio tracks generated from *P. cynomolgi* salivary gland sporozoites and blood stage parasites. “Blue genes” are transcribed from left to right, while “red genes” from right to left.

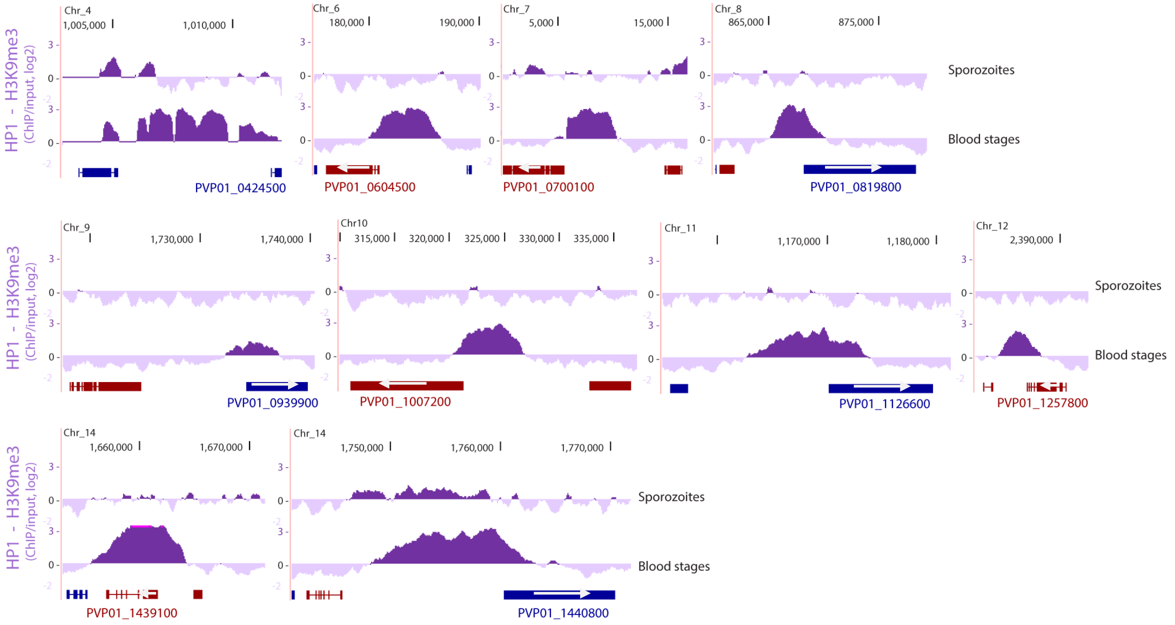


**Figure S1B: Epigenetic profile of *P vivax* genes listed on Fig. 1D**

Log2 transformed H3K9me3 - salivary gland sporozoites (upper, [16]) and HP1 - blood stage (lower, [15]) ChIP-over-input ratio tracks of *P. vivax* field isolates. “Blue genes” are transcribed from left to right, while “red genes” from right to left.

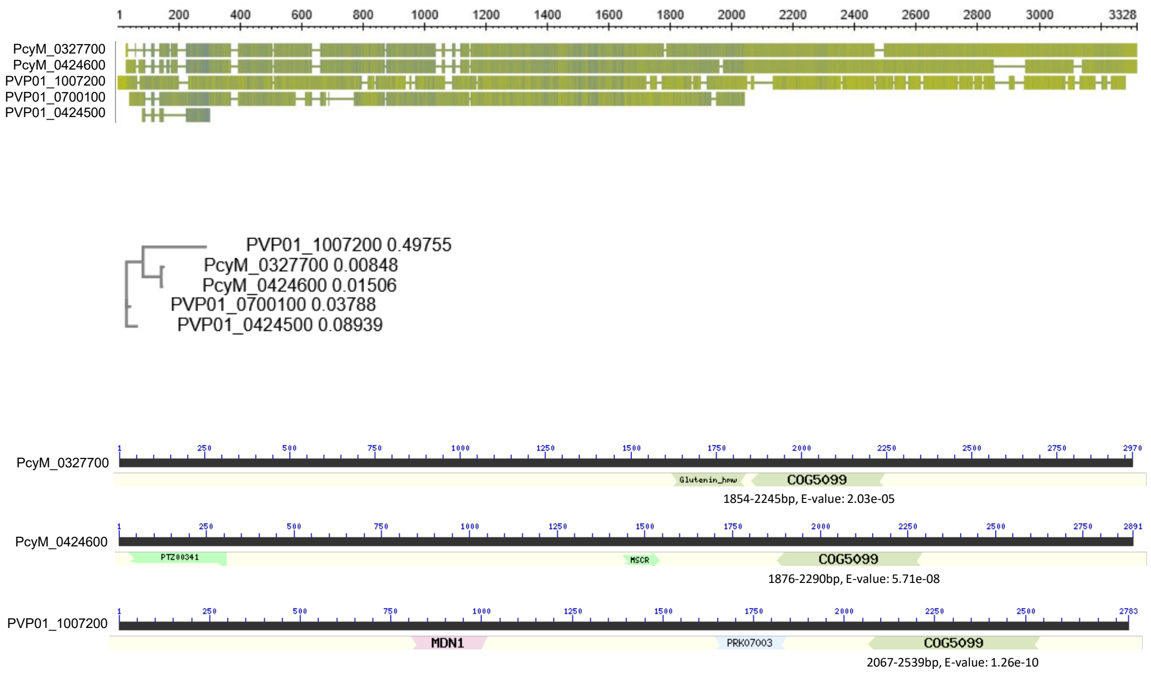


**Figure S1C: A small heterochromatin gaining gene family with Pumilio homology domain in *P. cynomolgi* and *P. vivax***

**Top:** multiple sequence alignment of two paralogues from *P. cynomolgi* and three paralogues from *P. vivax*. Note that two paralogues in *P. vivax* encode for substantially shorter / truncated proteins.

**Middle:** phylogenetic tree depicting the relatedness of the family members in *P. cynomolgi* and *P. vivax*. Note that the two *P. cynomolgi* paralogs are more similar to each other suggesting a “recent” gene duplication event.

**Bottom:** position of the Pumilio homology domain (COG5099) within the three full length paralogs as identified by NCBI conserved domain search. Note that other putative domains are less consistently identified across the paralogs.

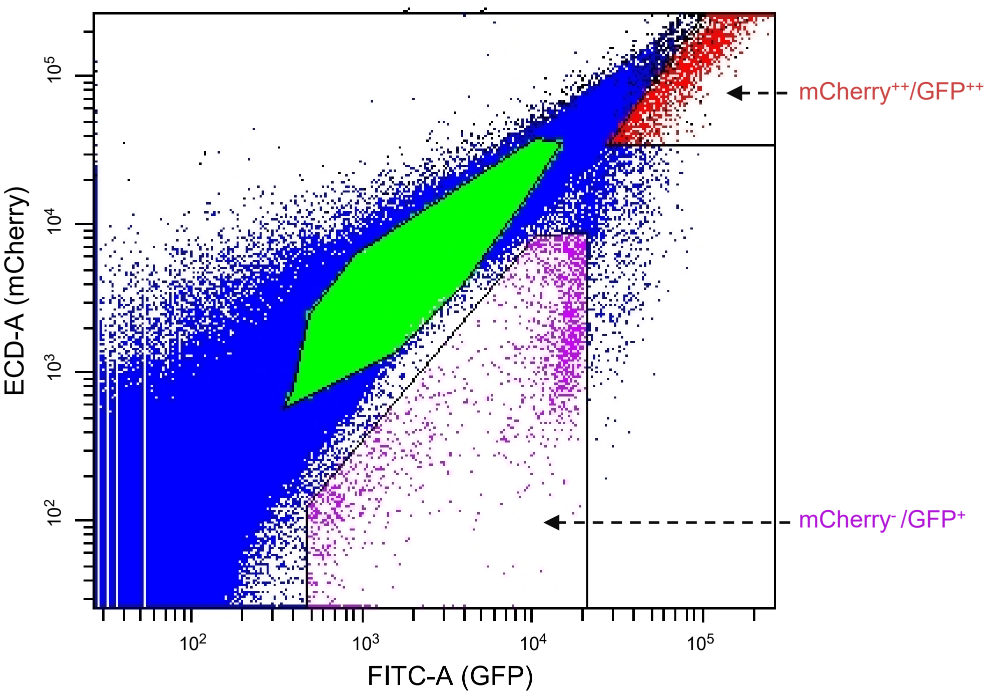


**Figure S2A: FACS gating strategy to isolate developing liver stage double-positive forms (mCherry++/GFP++) and small liver stage forms (mCherry-/GFP+) at 6 days post infection.**

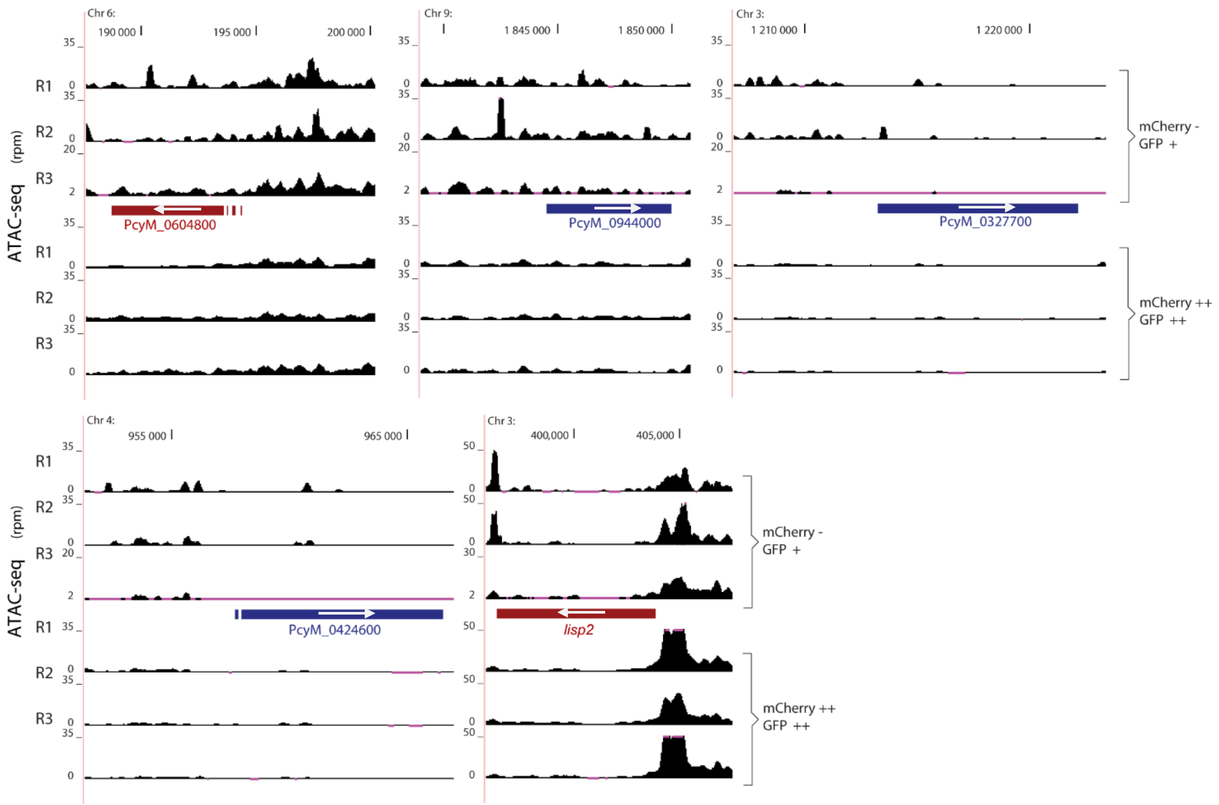


**Figure S2B: Chromatin accessibility profiles of *P. cynomolgi* genes named on Fig. 2B**
ATAC-seq read per million coverage tracks of mCherry-/GFP+ small liver stage forms and mCherry++/GFP++ liver schizonts over loci encoding for the heterochromatin gaining RNA-binding proteins (PcyM_0604800; PcyM_0944000; PcyM_0327700; PcyM_0424600) as well as *lisp2*. "Blue genes" are transcribed from left to right, while "red genes" from right to left.

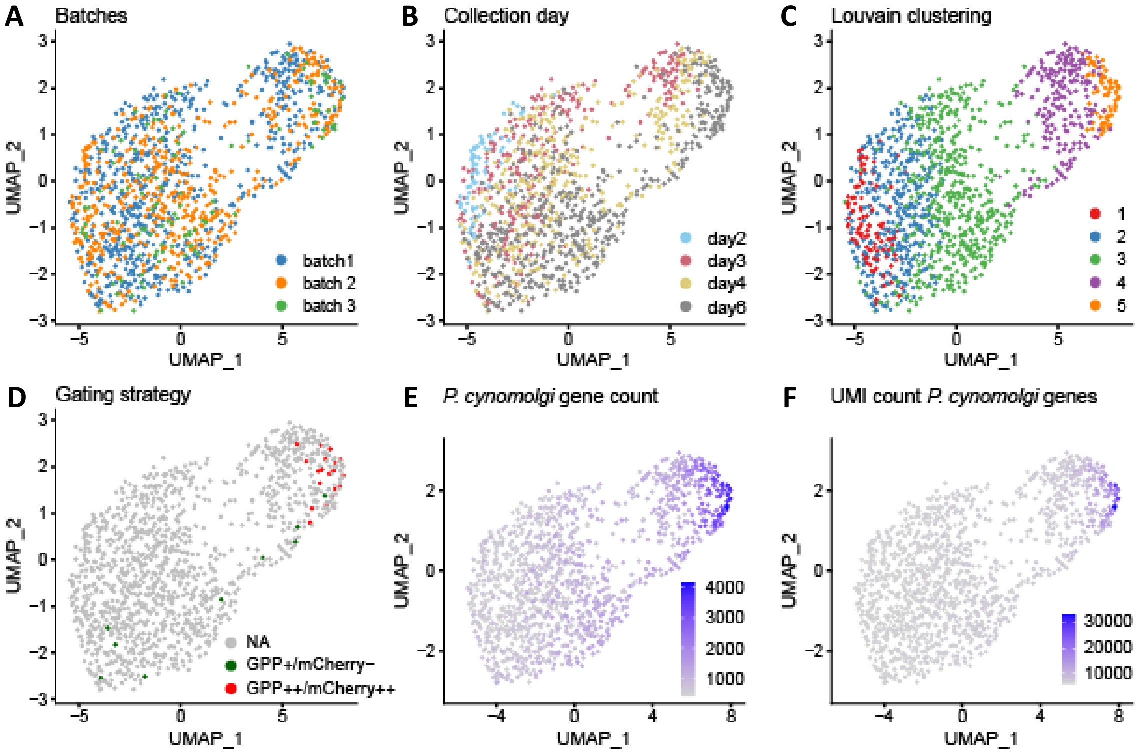


**Figure S3: Single-cell data analysis**

Uniform Manifold Approximation and Projection (UMAP) of 1243 liver stage *P. cynomolgi* parasites based on their gene expression profile. **A,** colored based in experimental batches; **B,** colored based on age of the parasites (days post infection); **C,** colored based on unsupervised clustering of expression profiles (Louvain clustering at resolution 0.4 of PCs 1-15). **D**, twenty index-sorted day 6 mCherry$^-$/GFP$^+$ (green) and mCherry$^{++}$/GFP$^{++}$ (red) parasites are highlighted; **E,** colored based on detected gene count; **F**, colored based on the number of unique molecular identifiers detected.

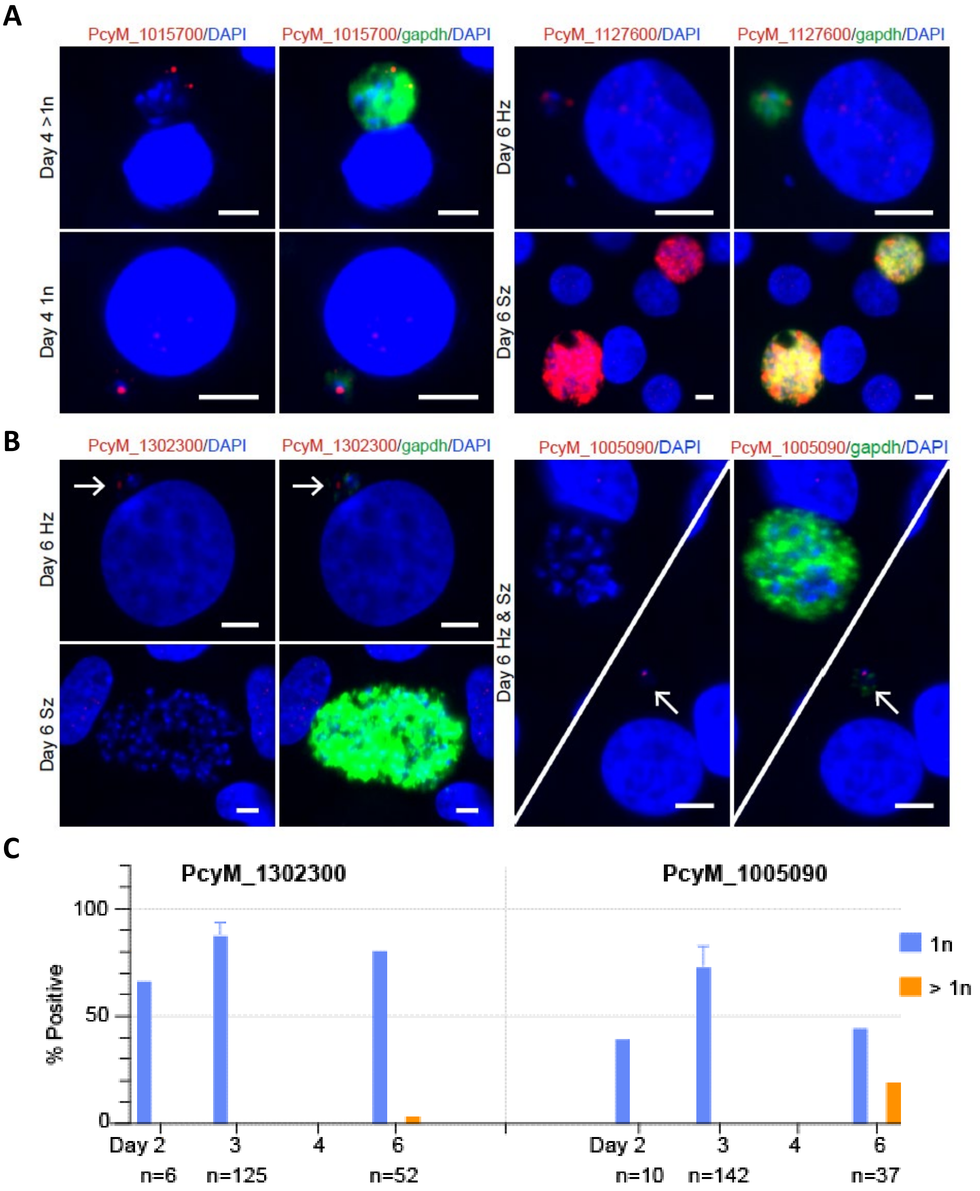


**Figure S4: FISH analysis of four *P. cynomolgi* genes**

**A**) Fluorescence *in situ* RNA hybridization images of two ApiAP2 transcription factor gene transcripts (PcyM_1015700 / *ap2-o2* and PcyM_1127600 / *ap2-sp3*, red) in relation to the house-keeping gene transcript (PcyM_1250000 / *gapdh*, green) in *P. cynomolgi* parasite-infected hepatocyte cultures.

**B**) Fluorescence *in situ* RNA hybridization images of two cluster II gene transcripts (PcyM_1302300 / *tgs1* and PcyM_1005090 / *conserved Plasmodium protein*, red) in relation to the house-keeping gene transcript (PcyM_1250000 / *gapdh*, green) in *P. cynomolgi* parasite-infected hepatocyte cultures. DNA of the parasites as well as host hepatocytes is visualized by DAPI staining (blue). Scale bar: 5 µm

**C**) Bar graph depicting the percentage of uninucleate (1n) and multinucleate (>1n) liver stage *P. cynomolgi* liver stage parasites with FISH signal using probes against PcyM_1302300 and PcyM_1005090 respectively at different days post infection. Error bars depict standard deviation.

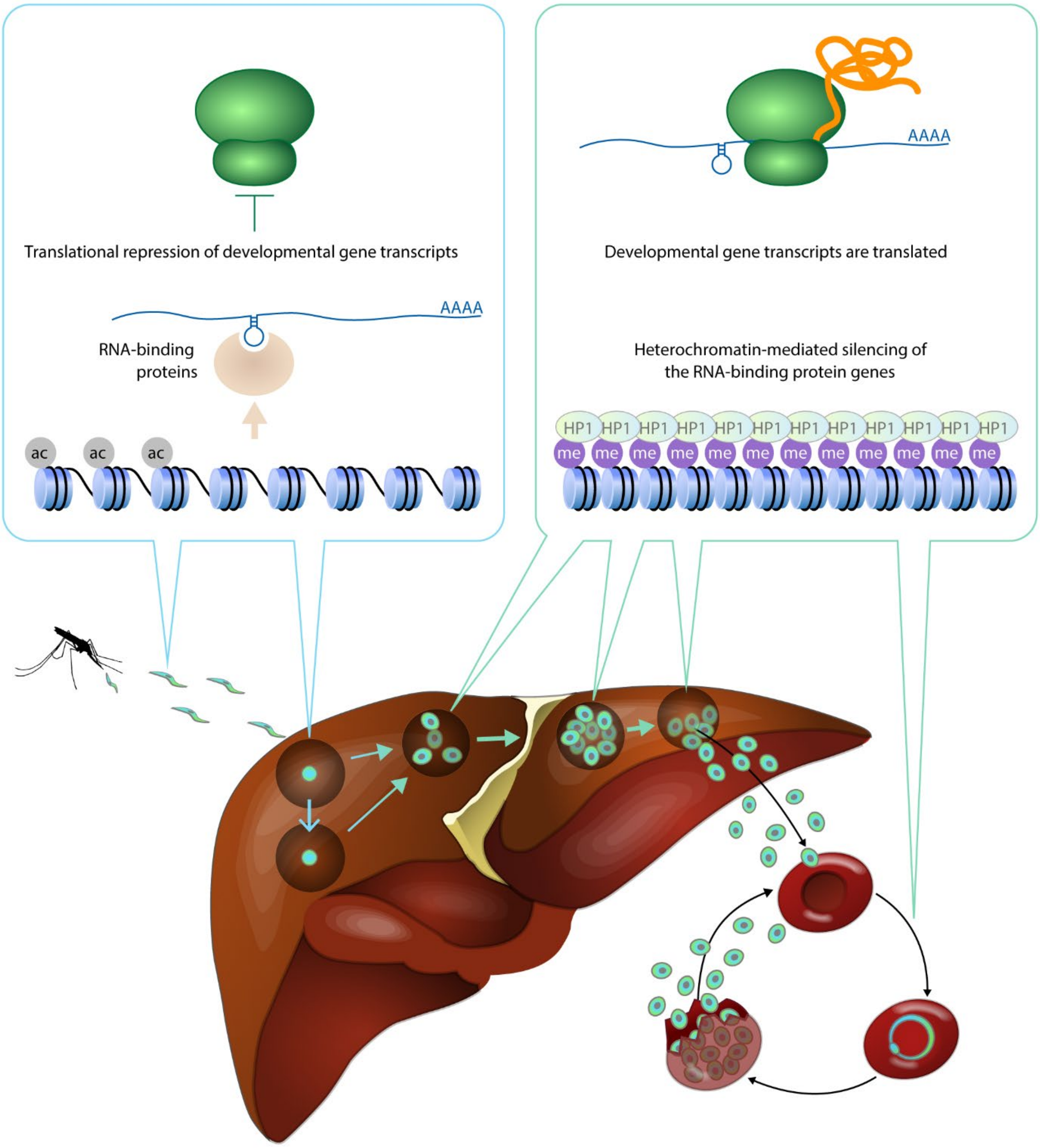


**Model: Hypothetical model of hypnozoite formation and reactivation**

Here we identified a small set of epigenetically-regulated RNA-binding proteins the are specifically expressed in small liver stage forms of relapsing malaria parasites leading to the following model. Genes encoding these RNA-binding proteins are in an euchromatic (i.e. acetylated) state in most, if not all sporozoites entering the liver. Given the euchromatic state of these genes they give rise to RNA-binding proteins, which in turn keep some transcripts essential for developmental progression in an unprocessed or translationally repressed state. Accordingly, in most small liver stage forms and in particular in hypnozoites development is halted. Upon *de novo* heterochromatin formation over the RNA-binding protein genes, these repressive proteins are not produced anymore and therefore developmental gene transcripts are efficiently translated, resulting in continuation of the developmental program (liver stage proliferation and/or hypnozoite reactivation).

| Gene ID | Description | Region targeted | Probe type |
|---|---|---|---|
| PcyM_1250000 | glyceraldehyde-3-phosphate dehydrogenase, putative | 113–997 | C1 |
| PcyM_0604800 | ZF-protein putative | 995-1987 | C2 |
| PcyM_0944000 | RNA-binding protein | 1651-2782 | C2 |
| PcyM_1302300 | trimethylguanosine synthase, putative | 28-1236 | C2 |
| PcyM_1015700 | AP2 domain transcription factor AP2-O2, putative | 1938-2904 | C2 |
| PcyM_1005090 | Conserved Plasmodium protein, unknown function | 2-423 | C2 |
| PcyM_1127600 | AP2 domain transcription factor AP2-SP3, putative | 8130-9313 | C2 |

**Table S5: RNAscope probes used for *in situ* fluorescence RNA hybridization**

<table>
<tr><th rowspan="2">GeneID</th><th rowspan="2">Predicted funcion</th><th rowspan="2">P. vivax</th><th colspan="2">Epigenetic regulation</th><th colspan="7">Hypnozoite-specific/enhanced expression</th></tr>
<tr><th>ChIP-seq</th><th>ATAC-seq</th><th>scRNA-seq</th><th>FISH</th><th>Cubi '17</th><th>Voorberg '17</th><th>Gural '18</th><th>M-Silva '22</th><th>Ruberto '22</th></tr>
<tr><td>PcyM_0604800</td><td>RNA-binding (ZF)</td><td>PVP01_0604500</td><td>Heterochromatin gaining</td><td>Differentialy accessible</td><td>+</td><td>+</td><td>+</td><td>+</td><td>+</td><td>+</td><td>-</td></tr>
<tr><td>PcyM_0944000</td><td>RNA-binding (RRM, BMS1)</td><td>PVP01_0939900</td><td>Moderately heterochr. gaining</td><td>Differentialy accessible</td><td>+</td><td>+</td><td>+</td><td>+</td><td>+</td><td>+</td><td>+</td></tr>
<tr><td>PcyM_0327700</td><td>RNA-binding (PUF)</td><td rowspan="2">PVP01_1007200</td><td>Heterochromatin gaining</td><td>Differentialy accessible</td><td>+/-</td><td>ND</td><td>+</td><td>+</td><td rowspan="2">+</td><td rowspan="2">+</td><td rowspan="2">+</td></tr>
<tr><td>PcyM_0424600</td><td>RNA-binding (PUF)</td><td>Heterochromatin gaining</td><td>Differentialy accessible</td><td>+/-</td><td>ND</td><td>-</td><td>-</td></tr>
<tr><td>PcyM_1302300</td><td>RNA methyltransferase</td><td>PVP01_1301800</td><td>Euchromatic</td><td>Not differentialy accessible</td><td>+</td><td>+</td><td>+</td><td>+</td><td>+</td><td>+/-</td><td>+/-</td></tr>
<tr><td></td><td></td><td></td><td></td><td></td><td></td><td></td><td></td><td></td><td></td><td></td><td></td></tr>
<tr><td>PcyM_1127600</td><td>DNA-binding (AP2-SP3)</td><td>PVP01_1126600</td><td>Heterochromatin gaining</td><td>Not differentialy accessible</td><td>+/-</td><td>-</td><td>+</td><td>+</td><td>-</td><td>+</td><td>+/-</td></tr>
<tr><td>PcyM_1015700</td><td>DNA-binding (AP2-O2)</td><td>PVP01_0734300</td><td>Euchromatic</td><td>Not differentialy accessible</td><td>+/-</td><td>-</td><td>+</td><td>-</td><td>-</td><td>-</td><td>+/-</td></tr>
<tr><td></td><td></td><td></td><td></td><td></td><td></td><td></td><td></td><td></td><td></td><td></td><td></td></tr>
<tr><td>PcyM_1005090</td><td>none predicted</td><td>PVP01_1007250</td><td>Euchromatic</td><td>Not differentialy accessible</td><td>+</td><td>+</td><td>-</td><td>-</td><td>+</td><td>-</td><td>-</td></tr>
</table>

**Table S6: Overview of the genes identified in this study being relevant for early liver stage development of relapsing malaria parasites and/or hypnozoite biology**
